# Comparative transcriptome profiling conferring of resistance to *Fusarium oxysporum* infection between resistant and susceptible tomato

**DOI:** 10.1101/116988

**Authors:** Min Zhao, Hui-Min Ji, Yin Gao, Xin-Xin Cao, Hui-Yin Mao, Peng Liu, Shou-Qiang Ouyang

**Affiliations:** College of Horticulture and Plant Protection; Texting Center; Joint International Research Laboratory of Agriculture and Agri-Product Safety; Key Laboratory of Plant Functional Genomics of the Ministry of Education, Yangzhou University, 48 East Wenhui Road, Yangzhou, Jiangsu, 225009, China

**Keywords:** Tomato, *Fusarium oxysporum* f. sp. *Lycopersici*, comparative transcriptome, pathogen, resistance

## Abstract

Tomato Fusarium wilt caused by *Fusarium oxysporum* f. sp. *lycopersici* (FOL) is a destructive disease of tomato worldwide which causes severe yield loss of the crops. As exploring gene expression and function approaches constitute an initial point for investigating pathogen-host interaction, we performed a transcriptional analysis to unravel regulated genes in tomato infected by FOL. Differentially expressed genes (DEG) upon inoculation with FOL were presented at twenty-four hours post-inoculation including four treatments: Moneymaker_H_2_O, Moneymaker_FOL, Motelle_H_2_O and Motelle_FOL. A total of more than 182.6 million high quality clean reads from the four libraries were obtained. A large overlap was found in DEGs between susceptible tomato cultivar Moneymaker and resistant tomato cultivar Motelle. All Gene Ontology terms were mainly classified into catalytic activity, metabolic process and binding. However, Gene Ontology enrichment analysis evidenced specific categories in infected Motelle. Statistics of pathway enrichment of DEGs resulted that the taurine and hypotaurine metabolism, the stibenoid, diarylheptanoid and gingerol biosynthesis, the starch and sucrose metabolism were the top three pathway affected in both groups. Interestingly, plant-pathogen pathway was greatly regulated in Motelle treated with FOL. Combining with qRT-PCR facilitated the identification of regulated pathogenicity associated genes upon infected resistant or susceptible tomato. Our data showed that a coordinated machinery played a critical role in prompting the response, which could help in generating models of mediated resistance responses with assessment of genomic gene expression patterns.

## INTRODUCTION

*Fusarium oxysporum* f. sp. *lycopersici* (hereafter referred to as FOL) is a biotrophic pathogen which is the causal agent of tomato wilt. Accumulating data indicate that *F. oxysporum* is a large species complex, with more than 150 host-specific forms causing disease in vegetables, fruit trees, wheat, corn, cotton and ornamental crops (Di Pietro *et al.* 2003; Leslie and Summerell 2006). F. *oxysporum* infects vascular bundles in the plant host, leading to clogged vessels, yellowing of leaves, wilting and finally death of the whole plant. According to their specific pathogenicity to tomato cultivars, three physiological races (Di Pietro *et al.* 2003; Leslie and Summerell 2006; Takken and Rep 2010) of *F. oxysporum* are distinguished (Kawabe *et al.* 2005).

Tomato *(Solanum lycopersicum)* is a worldwide economic crop, and also has been studied as a crucial model plant for studying the genetics and molecular basis of resistance mechanisms. Four plant resistance *(R)* genes have been discovered in cultivated tomato from wild tomato species including the *I* and *I-2* genes from *S. pimpinellifolium,* and the *I-3* and *I-7* gene from *S. pennellii.* Among these four *R* genes, so far, *I-2, I-3* and *I-7* have been cloned, encode an NB-LRR protein like most known *R* genes (Ori *et al.* 1997; Simons *et al.* 1998; Kawabe *et al.* 2005; Catanzariti *et al.* 2015; Gonzalez-Cendales *et al.* 2016). Previous works have demonstrated that the *I-2* and *I-3* gene confers resistance to race 2 and race 3 strains of FOL, respectively (Simons *et al.* 1998; Catanzariti *et al.* 2015). The *I-2* locus encodes an *R* protein that recognizes the *avr2* gene product from *F. oxysporum* (race 2) (Houterman *et al.* 2009). The *I-3* encodes an S-receptor-like kinase (SRLK) genes that confers Avr3-dependent resistance to FOL (race 3) (Catanzariti *et al.* 2015). Previously, two near-isogenic tomato cultivars susceptible Moneymaker *(i-2/i-2)* and resistant Motelle (I-2/I-2) were recruited to study the interaction between tomato and FOL (Ouyang *et al.* 2014). The genotypes of these two tomato cultivars are for *I-2* and respond to FOL infection (Di Pietro and Roncero 1998; De Ilarduya *et al.* 2001; Yu and Zou 2008). We unrevealed the microRNA diversifications responding to FOL infection in tomato by high-throughput RNA sequencing (RNA-seq) approach (Ouyang *et al.* 2014). Basically, transcriptome analysis is a very important tool to discover the molecular basis of plant-pathogen interaction globally, allowing dissection of the pattern of pathogen activities and molecular repertoires available for defense responses in host plant. By taking advantage of RNA-seq technology, a few of transcriptome profiling studies of plants following inoculation with *Fusarium* fungus have been reported, including studies in banana (Guo *et al.* 2014), cabbage (Xing *et al.* 2016), watermelon (Liu *et al.* 2015), mango (Liu *et al.* 2016), and *Arabidopsis* (Chen et al. 2014; Gupta *et al.* 2014). Upon to pathogens infection, plants activate a few of defense responses to resistant diseases caused by according pathogens. Resistance response may associated with hypersensitive reaction (HR), structural alterations, reactive oxygen species (ROS) accumulation, synthesis of secondary metabolites and defense molecules (Park *et al.* 2003; Shah 2003; Ros *et al.* 2004).

The objects of this study were to determine the transcript profile between susceptible Moneymaker and resistant Motelle tomato plants in response to FOL infection and to reveal genes underlying the innate immune response against the fungal pathogen. To achieve these goals, we performed transcriptome analysis using RNA-seq approach. In addition to genes known to response to pathogen infection, our results also uncovered a bunch of novel fungal pathogen-responsive genes for further functional characterization, and provided a broader view of the dynamics of tomato defense transcriptome triggered by FOL infection.

## MATERIALS and METHODS

### Tomato materials and fungal culture

Two tomato near-isogenic cultivars (cv.) Motelle *(I-2/I-2)* and Moneymaker *(i-2/i-*2) that exhibit different susceptibilities to the root pathogen FOL were used for plant infection and libraries construction. Profiling experiments were performed on two-week-old tomato seedlings grown at 25°C with a 16/8-h light/dark cycle. The wild-type *Fusarium oxysporum* f. sp *lycopersici* strain used for all experiments is FGSC 9935 (also referred to as FOL 4287 or NRRL 34936). Two-week-old tomato seedlings were removed from soil and roots incubated in a solution of FOL conidia at a concentration of 1x10^8^/ml for 30 min. Control tomato plants were treated with water. Plants were then replanted in soil and maintained in a growth chamber at 25°C for 24 h with constant light. Plants were removed from soil, and roots were rinsed and excised, then immediately frozen in liquid nitrogen and stored at −80°C.

### RNA extraction, library preparation, and sequencing

Total RNA was isolated from roots using TRIzol^®^ Reagent (#15596026, Life Technologies, CA, USA) according to the manufacturer’s recommendations. After the total RNA extraction and DNase I treatment, magnetic beads with Oligo (dT) were used to isolate mRNA. Mixed with the fragmentation buffer, the mRNA was sheared into short fragments. Then cDNA was synthesized using the mRNA fragments as templates. cDNAs were purified and resolved with EB buffer for end reparation and single nucleotide A (adenine) addition followed by adding adapters to cDNAs. After agarose gel electrophoresis, the suitable cDNAs were selected for the PCR amplification as templates. During the quality control (QC) steps, Agilent 2100 Bioanaylzer and ABI StepOnePlus Real-Time PCR System were used in quantification and qualification of the sample library. The libraries were sequenced using Illumina HiSeqTM 2000.

### RNA-seq analysis, normalization of sequence reads and identification of differentially expressed genes (DEGs)

Primary sequencing data that produced by Illumina HiSeqTM 2000, called as raw reads, were subjected to QC. After QC, raw reads were filtered into clean reads which were aligned to the reference sequences as described by previous report. (Trapnell *et al.* 2012). All sequence reads were trimmed to remove the low-quality sequences. The trimmed reads were then aligned to the tomato reference genome downloaded from the Sol Genomics Network using Bowtie v0.12.5 (Langmead *et al.* 2009) and TopHat v2.0.0 (Trapnell *et al.* 2009; Trapnell *et al.* 2012) with default settings. Cufflinksv0.9.3 (Trapnell *et al.* 2010) was used to calculate transcript abundance based on fragments per kilo base of transcript permillion fragments mapped (FPKM) using all parameters on default settings. The transcript was considered as expressed when the FPKM value was greater than 0.1 and the lower boundary for FPKM value was greater than zero at 95% confidence interval. Once the transcript abundance was calculated for individual sample files using Cufflinks, the output files were further merged pairwise for each comparison (in vitro comparison between two populations, in planta comparison between two populations and in planta versus *in vitro* for each population) using Cufflinks utility program-Cuffmerge (Trapnell *et al.* 2012). The pairwise comparisons of gene expression profiles between the two populations were done using the Cuffdiff program of the Cufflinks version 1.3.0 (Trapnell *et al.* 2010). The genes were considered significantly differentially expressed if Log_2_ FPKM (fold change) was ≥1.0 and false discovery rate (FDR, the adjusted P value) was <0.01. The q-value which was a positive FDR analogue of the p-value was set to <0.01 (Storey and Tibshirani 2003).

### Functional categorization of DEGs

DEGs were functionally categorized online for all pairwise comparisons according to the Munich Information Center for Protein Sequences (MIPS) functional catalogue (Ruepp *et al.* 2004). The functional categories and subcategories were regarded as enriched in the genome if an enrichment P- and FDR-value was below <0.05. The Kyoto Encyclopediaof Genes and Genomes (KEGG) pathway analyses were performed using interface on blast2GO (Blast2GO v2.6.0, http://www.blast2go.com/b2ghome) for all DEGs to identify gene enrichment on a specific pathway.

### Gene Ontology (GO) and pathway enrichment analysis

Gene Ontology (GO) and pathway enrichment were performed using DAVID software (Smyth 2005). Graphs of the top 20 enriched GO terms for each library were generated using the Cytoscape Enrichment Map plugin (Smoot *et al.* 2011; Merico *et al.* 2010).

### Quantitative real time-PCR (qRT-PCR) analysis

qRT-PCR analysis was performed according to our previous protocol (Ouyang et al. 2014). The reverse transcription reaction was done on 1 μg of total RNA using the SMART MMLV Reverse Transcriptase (Takara, Mountain View, CA). cDNA was diluted two times and used as template for quantitative RT-PCR, which was performed with the CFX96 real-time PCR system (Bio-Rad, Hercules, California, USA). Primers used for qRT-PCR were designed from 3-UTR for individual gene. For each cDNA sample, three replications were performed. Each reaction mixture (20 μL) contained 1 μL of cDNA template, 10 μL of SYBR1 Green PCR Master Mix (Applied Biosystems, Foster, CA) and 1 μL of each primer (10 μM). Relative expression levels of genes were normalized using the 18S rRNA as internal control, and were calculated as the fold change by comparison between in water treated and in FOL treated samples.

### Statistical analyses

All data in this study were subjected to ANOVA analysis or Student’s t-test analysis using SPSS 11.5 (SPSS Company, Chicago, IL).

## RESULTS

### General features of Moneymaker and Motelle transcriptomes

We investigated transcriptomes in roots of tomato during infection with the tomato wilt disease fungus FOL through construction of transcript libraries and RNA-seq. By taking advantage of two near-isogenic cultivars that show differential interaction with FOL - Moneymaker (susceptible) and Motelle (resistant), we generated four libraries including: Moneymaker treated with water (MM_H_2_O), Moneymaker treated with FOL (MM_Foxy), Motelle treated with water (Mot_H_2_O) and Motelle treated with FOL (Mot_Foxy). Using Illumina sequencing, we obtained a total of more than 182.6 million high quality clean reads from the four libraries. Of these, 45,616,330 from MM_H_2_O, 45,635,428 from MM_Foxy, 45,680,034 from Mot_H_2_O, and 45,661,734 from Mot_Foxy. The number of expressed transcripts were 22,796 and 22,639 from MM_H_2_O and MM_Foxy library respectively, and 22,825 and 22,725 from Mot _H_2_O and Mot _Foxy library respectively (Table 1). Sequence reads presented reasonable correlation between related two populations (t >0.95, p = 0.29) (Figure 1). A number of 21,808 and 21,753 genes were co-expressed between MM_H_2_O and MM_Foxy library and Mot _H_2_O and Mot _Foxy library, respectively. The co-expressed genes increased slightly to 21,887 between MM_Foxy and Mot _Foxy library (Figure 2). The scatter of all expressed genes of each pair were presented in figure 3.

**Table 1.** Summary of sequence reads (in millions) from four libraries.

| Sample name | Clean reads | Genome map rate | Gene map rate | Expressed gene | Novel gene | Alternative splicing | SNP | Indel |
| --- | --- | --- | --- | --- | --- | --- | --- | --- |
| MM_H<sub>2</sub>O | 45,616,330 | 75.49% | 76.87% | 22,796 | 775 | 32,482 | 14,046 | 1,232 |
| MM_Foxy | 45,635,428 | 67.89% | 68.47% | 22,639 | 761 | 32,689 | 13,371 | 1,228 |
| Mot_H<sub>2</sub>O | 45,680,034 | 75.87% | 75.92% | 22,825 | 795 | 33,706 | 17,286 | 1,506 |
| Mot_Foxy | 45,661,734 | 70.46% | 71.26% | 22,725 | 705 | 32,965 | 16,409 | 1,360 |

**Figure 1.**
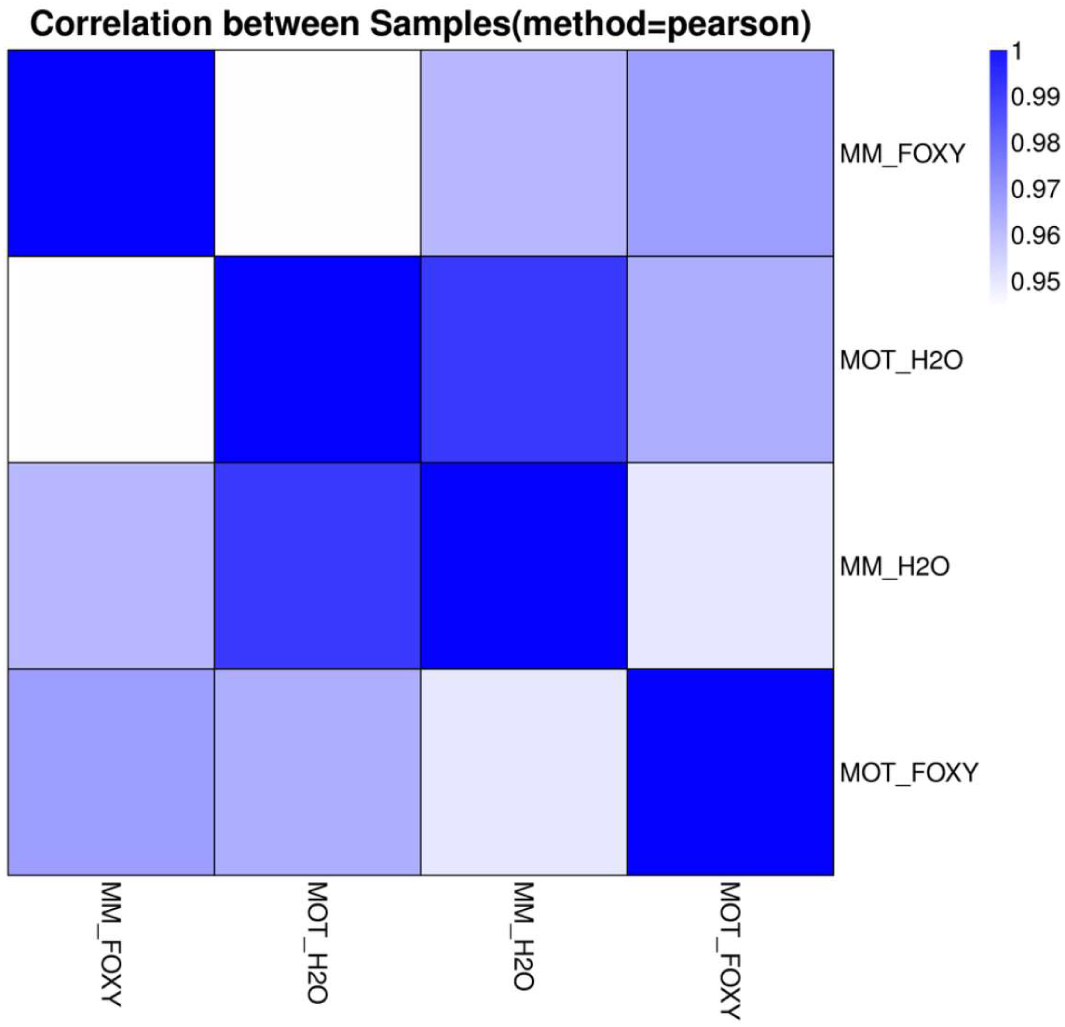
Correlations value between each two libraries.

**Figure 2.**
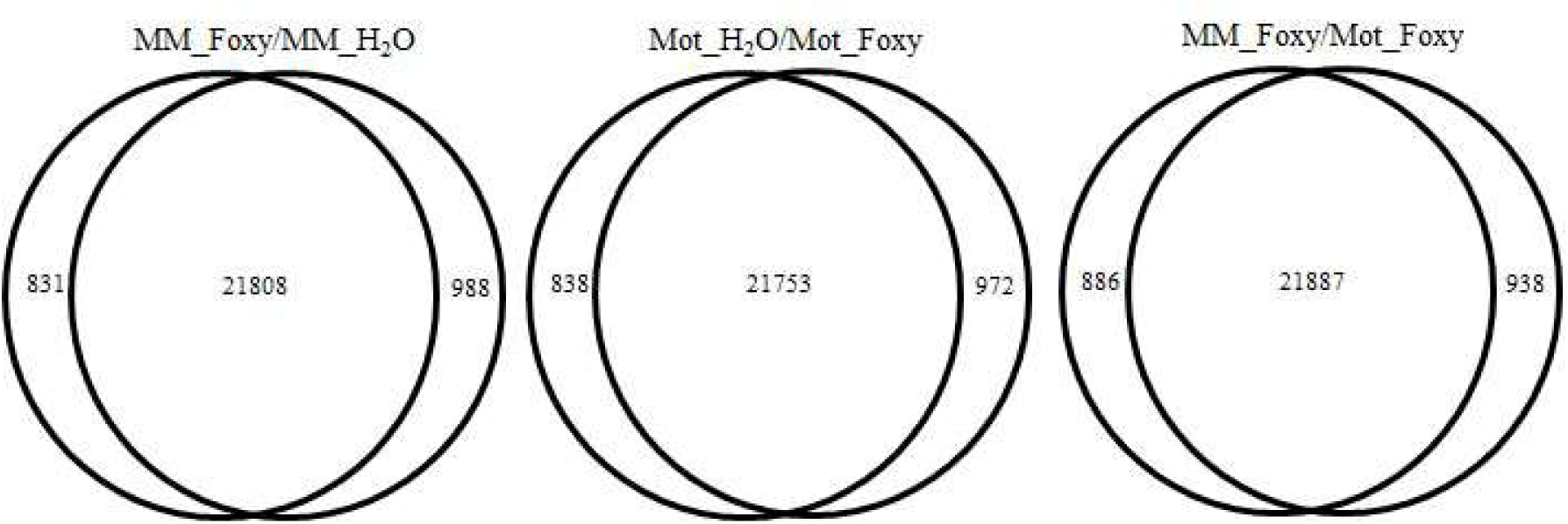
Venn Chart of Co-expressed Genes between MM_H_2_O and MM_Foxy, Mot _H_2_O and Mot _Foxy, and MM_Foxy and MM_Foxy.

**Figure 3.**
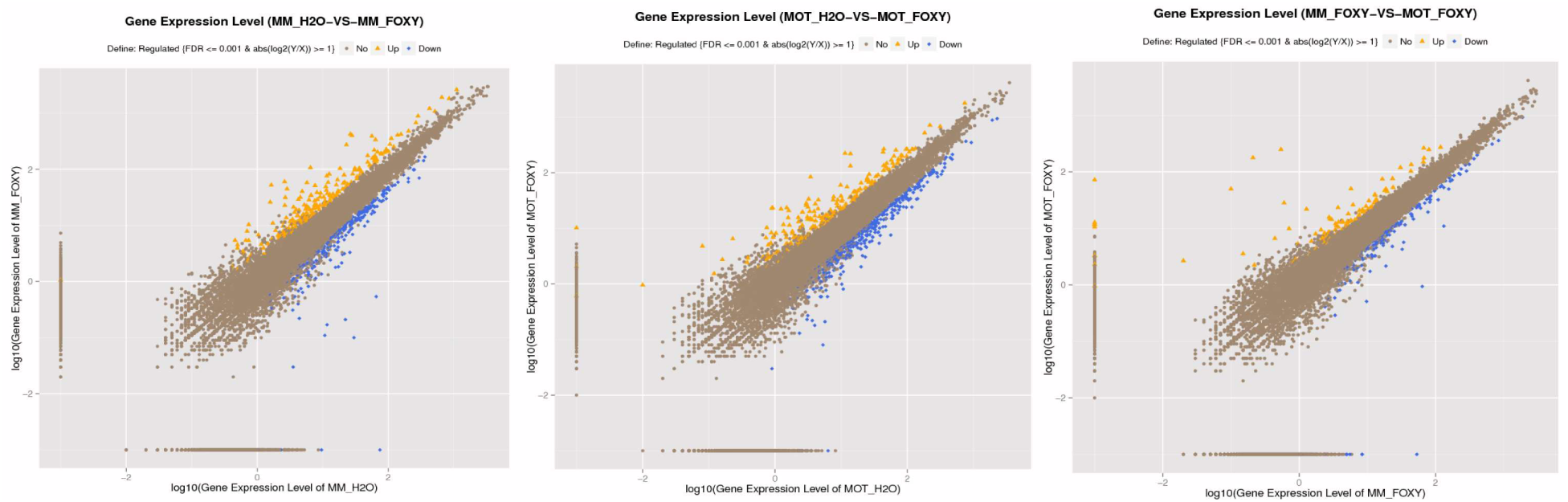
Scatter chart of all expressed genes of each pair between MM_H_2_O and MM_Foxy, Mot _H_2_O and Mot _Foxy, and MM_Foxy and MM_Foxy.

Of the sequence reads from MM_H_2_O and MM_Foxy library, 75.49% and 67.89% were mapped to the reference genome of tomato, respectively. For Mot _H2O and Mot _Foxy library, 75.87% and 70.46% were aligned to the reference genome of tomato, respectively. Among the reads mapped to the tomato genome, perfect match reads were 63.00% and 55.52% for MM_H_2_O and MM_Foxy library respectively, and 62.61% and 57.41% for Mot _H_2_O and Mot _Foxy library respectively (Table 2).

**Table 2.**
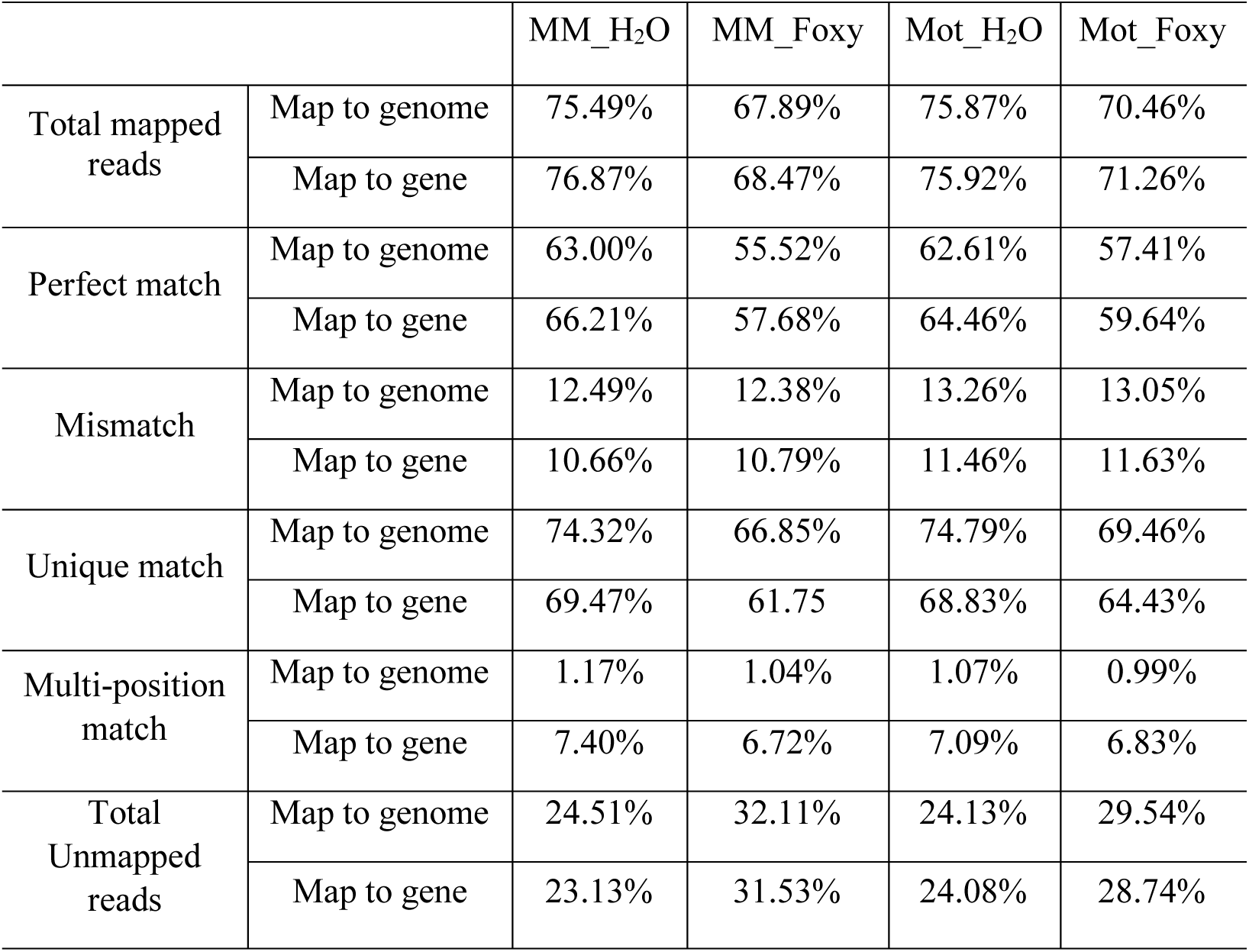
Mapping statistics of four libraries.

### Analysis of differentially expressed genes (DEGs) and functional classification of DEGs by gene ontology (GO) enrichment analysis

After expression levels Fragments Per Kilobase of exon model per Million mapped reads (FPKM) for each gene were calculated. Differentially expressed genes (DEGs) were defined as genes with fold-change > 2 fold and P_adjust_ value < 0.05. A total number of 3,942 and 4,168 genes showed significantly differential expression in MM_H_2_O vs. MM_Foxy library and Mot _H_2_O vs. Mot _Foxy library, respectively.

Among these DEGs, 221/219 genes were down-regulated, and 261/415 genes were up-regulated (MM_H_2_O vs. MM_Foxy/ Mot _H_2_O vs. Mot _Foxy) (Figure 4). A majority of these DEGs were overlapped in both water and FOL treated two tomato cultivars.

**Figure 4.**
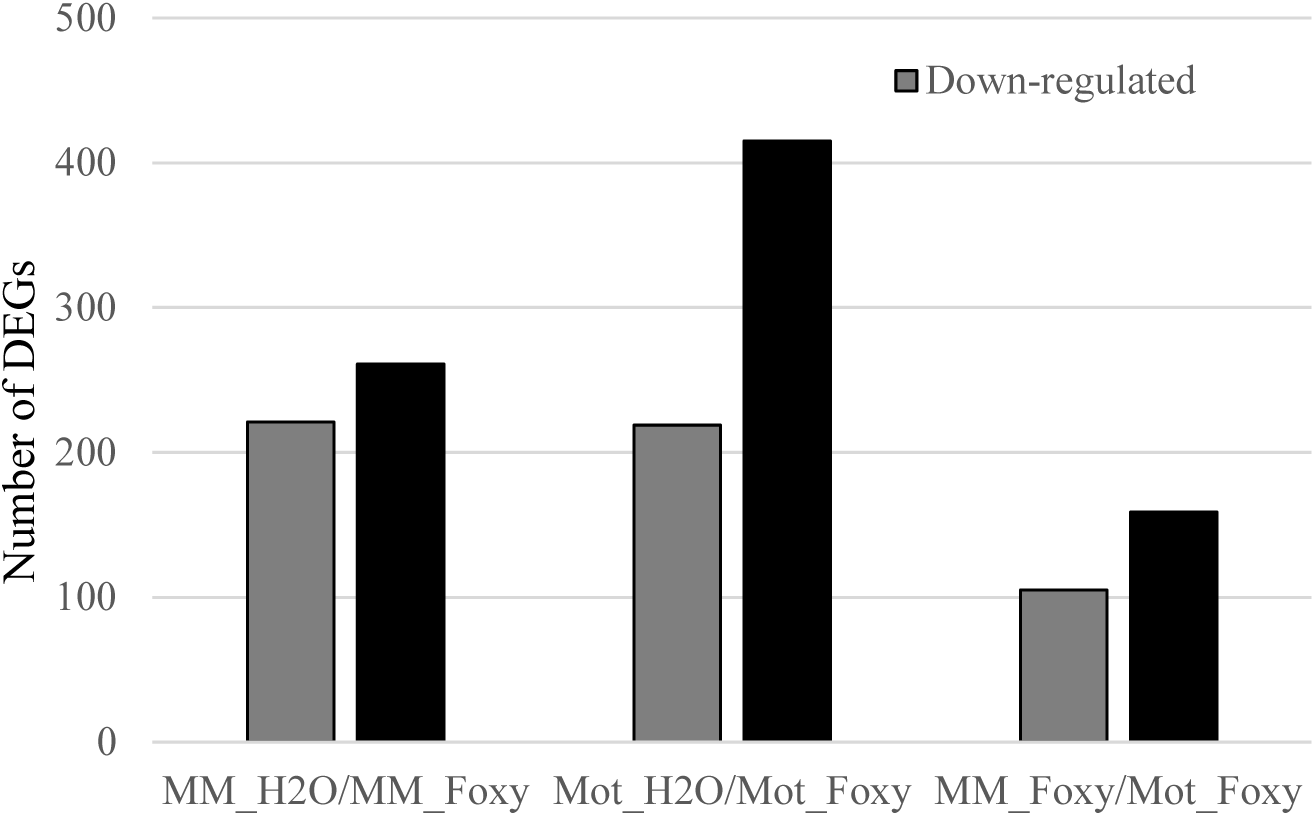
Statistics of Differentially Expressed Genes (DEGs).

To explore the distribution of DEGs, gene ontology (GO) enrichment analyses were conducted based on these DEGs. A total of 530 and 769 GO terms were discovered in MM_H_2_O vs. MM_Foxy and Mot _H_2_O vs. Mot _Foxy library, respectively. For both libraries, all GO terms were assigned to three groups including the biological process, the cellular component and the molecular. All GO terms were mainly classified into catalytic activity (104 out of 530 in MM_H_2_O vs. MM_Foxy library, and 141 out of 769 in Mot _H_2_O vs. Mot _Foxy library, (the same define in the following text), metabolic process (81 out of 530, and 118 out of 769), and binding (72 out of 530, and 104 out of 769). For the class of response to stimulus, however, no significant change was presented between these two libraries (31 out of 530, and 36 out of 769) (Figure 5).

**Figure 5.**
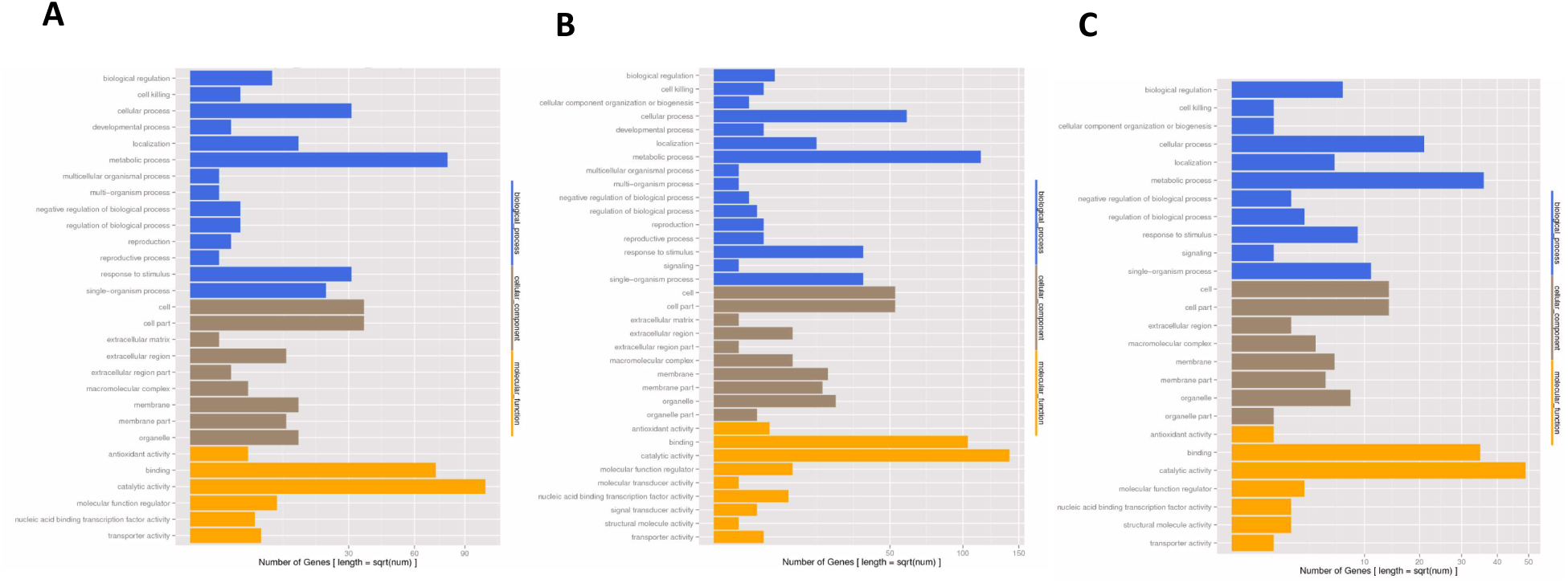
Gene Ontology Analysis of DEGs. The results were basically summarized into three main categories: biological processes, cellular components, and molecular functions. All statistically significant genes from four libraries were assigned to GO terms. A MM_H2O vs MM_Foxy. B Mot_H2O vs Mot_Foxy. C MM_Foxy vs Mot_Foxy.

To further understand the biological functions, top 20 statistics of pathway enrichment of DEGs were performed to discover the affection of FOL to host plant. Total of 356 and 469 DEGs from MM_H_2_O vs. MM_Foxy library and Mot _H_2_O vs. Mot _Foxy library respective were annotated for pathway enrichment. The taurine and hypotaurine metabolism, the stibenoid, diarylheptanoid and gingerol biosynthesis, the starch and sucrose metabolism were the top three pathway affected in both groups, but in different ranking. The metabolic pathway was the most abundant DEGs in both groups with 122 out of 356 and 148 out of 469 DEGs in MM_H_2_O vs. MM_Foxy library and Mot _H_2_O vs. Mot _Foxy library, respectively (Figure 6). Be worth mentioning, plant-pathogen pathway was ranked in the 24^th^ (24 out of 356 DEGs) in MM_H_2_O vs. MM_Foxy library, however, it was presented in the 8^th^ (40 out of 469 DEGs) in Mot _H_2_O vs. Mot _Foxy library (Figure 6). When compared with Mot _H_2_O vs. Mot_Foxy library, 19 DEGs were presented in MM_H_2_O vs. MM_Foxy library.

**Figure 6.**
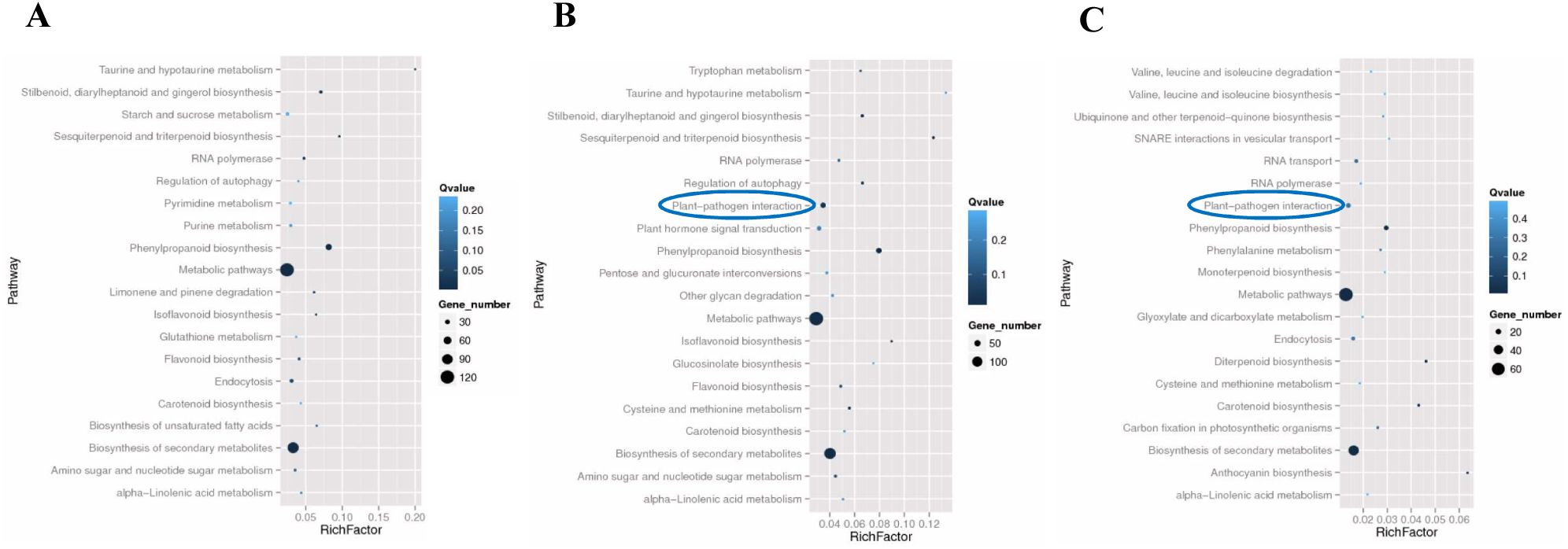
Top 20 of pathway enrichment analysis of DEGs. A MM_H2O vs MM_Foxy. B Mot_H2O vs Mot_Foxy. C MM_Foxy vs Mot_Foxy. Blue circle in B and C highlighted the plant-pathogen interaction.

### Expression profiles of DEGs selected in plant-pathogen interaction by qRT-PCR

To verify the DEGs in plant-pathogen interaction pathway, ten disease related DEGs were selected to characterize the gene expression profiles between water and FOL treated Moneymaker and Motelle by qRT-PCR using primers listed in table 3. These DEGs were Solyc00g174330 (Pathogenesis related protein PR-1), Solyc09g007010 (Pathogenesis related protein PR-1), Solyc02g084890 (Cc-nbs-lrr, resistance protein), Solyc07g054120 (LRR receptor-like serine/threonine-protein kinase, RLP), Solyc10g011910 (WRKY transcription factor 23), Solyc03g124110 (Pathogenesis-related transcriptional factor and ERF, DNA-binding), Solyc03g026280 (Pathogenesis-related transcriptional factor and ERF, DNA-binding), Solyc12g009240 (Pathogenesis-related transcriptional factor and ERF, DNA-binding) and Solyc02g080070 (RLK, Receptor like protein, putative resistance protein with an antifungal domain).

**Table 3.**
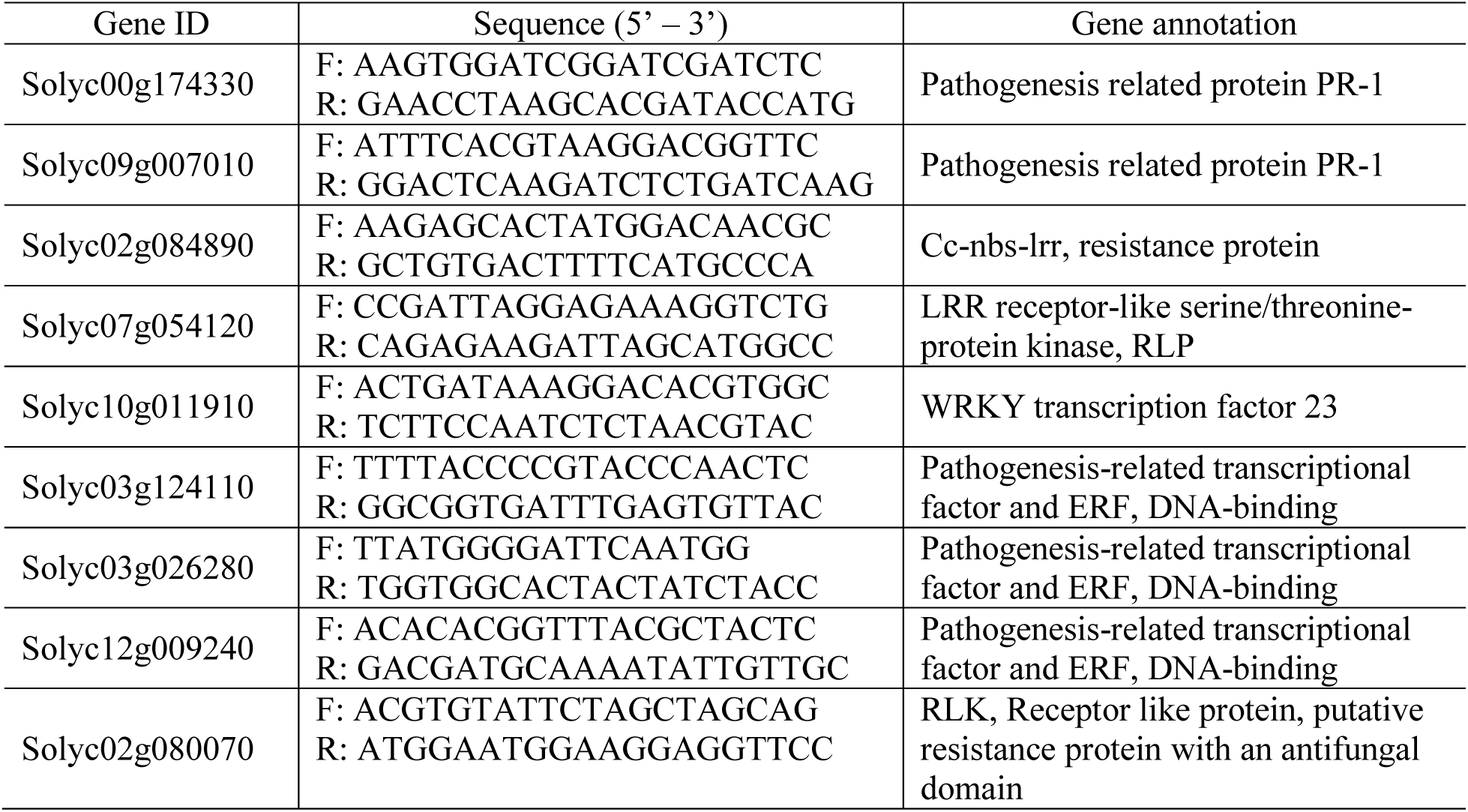
Annotation of pathogenesis related genes and primers used for qRT-PCR in this study

The results of qRT-PCR showed the similar pattern with sequencing results with minute difference. Among these DEGs, Solyc00g174330, Solyc10g011910, Solyc03g124110, Solyc02g084890 and Solyc12g009240 were induced greatly in Motelle affected by FOL, however, no significant changes were present in Moneymaker between water and FOL treatment (Figure 7).

**Figure 7.**
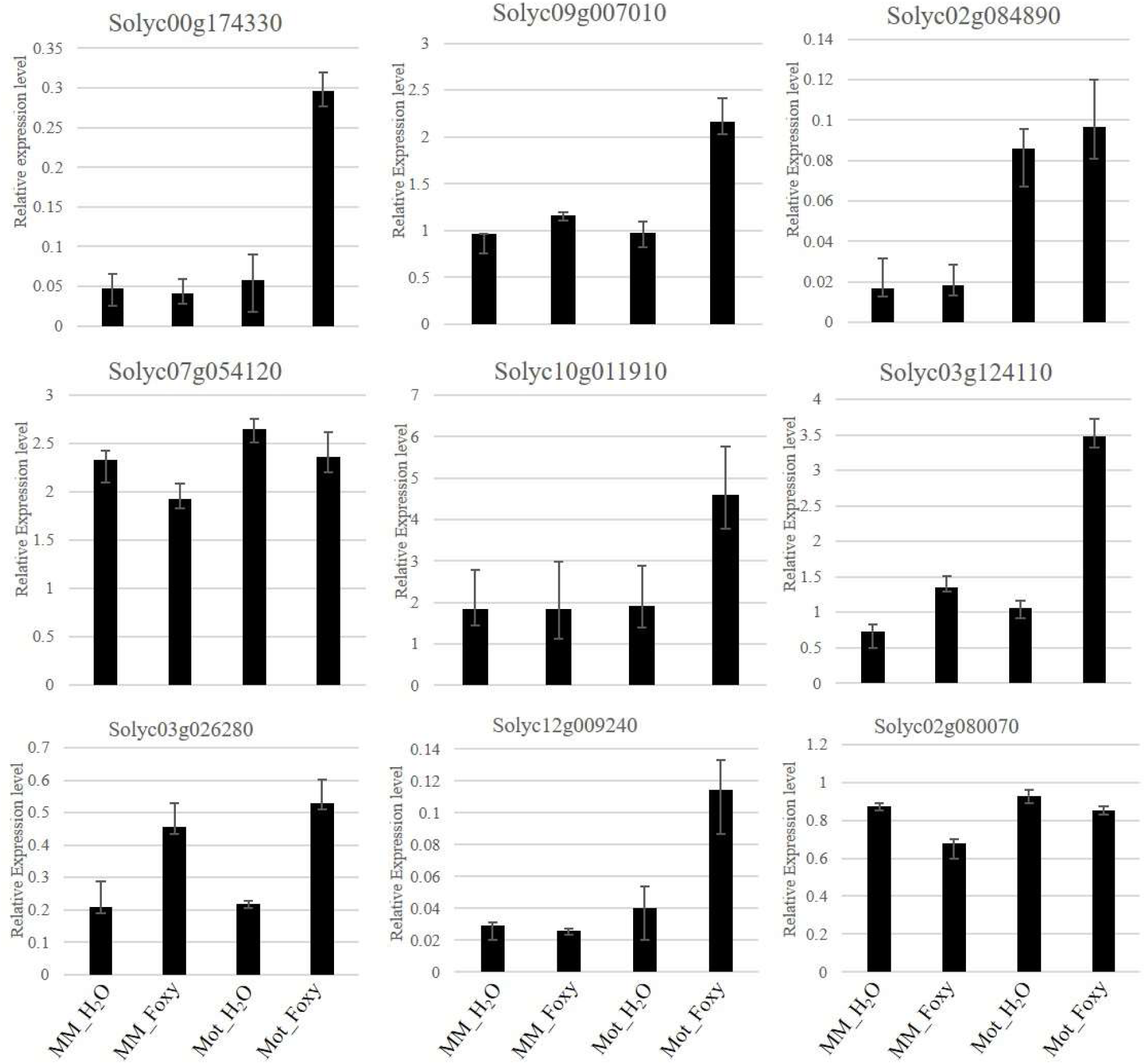
Validation of DEGs selected in plant-pathogen interaction pathway by qRT-PCR. Total tomato root RNA was used for qRT-PCR with gene-specific primers. Each column represents an average of three replicates, and error bars represent the standard error of means.

## DISCUSSION

In this study, we explored the availability of near-isogenic susceptible and resistant cultivars of tomato infected by FOL to uncover a global transcriptomic profile of tomato-FOL interaction using Illumina sequencing. The components of plant responding to pathogen challenging may lead to understand the underlying defense mechanisms. Plants have evolved a complicate defense system against pathogens including cascade signaling activation, the regulation of gene expression, synthesis of defensive metabolites as well as hormone balancing (Mukhtar *et al.* 2011; Andolfo *et al.* 2014). So far, by taking advantage of high-throughput RNAsequencing (RNA-seq) approach, a few of transcriptome studies discovering the *F. oxysporum-host* interaction have been reported in plants such as banana, watermelon, mango and Arabidopsis (Chen *et al.* 2014; Guo *et al.* 2014; Gupta *et al.* 2014; Liu *et al.* 2015; Liu *et al.* 2016; Xing *et al.* 2016), shedding light on the crosstalking among different signaling pathways involving in plant-pathogen interaction.

When plant is attacked by pathogen, the host reprograms metabolism balance between development and the resources to support defense to pathogen, involving biological process, cellular components and molecular functions (Mithöfer and Boland 2012). Upon to our results, the tomato-FOL interaction basically followed the typical reaction of biotrophic phase pathogens infection. Gene Ontology analysis of DEGs between two tomato cultivars revealed specific enriched categories in both interactions. In resistant tomato cultivar Motelle, cellular component organization or biogenesis, signaling, molecular transducer activity, and signal transducer activity were evidenced when compared to susceptive tomato cultivar Moneymaker. Among them, cellular component organization or biogenesis was a critical metabolic activities required by plants to survive under fungus-inflicted stresses (Paul *et al.* 2011). Generally, the genes involved in GO analysis presented in Motelle more than in Moneymaker upon FOL infection which was due to different resistant cultivar.

Two main mechanisms, pathogen-associated molecular patterns (PAMPs) or microbe-associated molecular patterns (MAMPs) (Boller and Felix 2009; Cui *et al.* 2014; Yang and Huang 2014) and the adaptive immune system composed of resistance (R) genes (Dangl and Jones 2001; Van Ooijen *et al.* 2007; Marone *et al.* 2013), are involved in plant responses to pathogenic microorganisms in plant. At least five different classes of *R* genes have been classified based on functional domain (Van Ooijen *et al.* 2007). Among these classes, a nucleotide-binding site (NBS) and leucine-reach repeats (LRRs) (NBS-LRR) is known as the most numerous R-gene class (Dangl and Jones 2001). Previously, we reported that tomato endogenic microRNA slmiR482f and slmiR5300 conferred to tomato wilt disease resistance. Two predicted mRNA targets each of slmiR482f and slmiR5300, encoded protein with full or partial NBS domains respectively, confirmed to exhibit function of resistance to FOL (Ouyang *et al.* 2014). A few of investments have been demonstrated that NB-LRR proteins are required for the recognition of a specific Avr and disease resistance in several plant species, including rice, *N. benthamiana,* Arabidopsis and wheat (Sinapidou *et al.* 2004; Peart *et al.* 2005; Lee *et al.* 2009; Loutre *et al.* 2009; Narusaka *et al.* 2009; Okuyama *et al.* 2011; Ouyang *et al.* 2014). The corresponding *R* genes were located tightly in physical linkage. However, in spite this physical linkage, not all these *R* gene pairs were homologous (Sinapidou *et al.* 2004; Lee *et al.* 2009). We found that genes related to plant-pathogen interaction were activated in resistant cultivar Moltelle once treated with FOL. Our qRT-PCR results demonstrated that some of these genes were up-regulated specifically in Motelle but not in Moneymaker. In particular, most of these genes were NBS-LRR or like genes which may imply that NBS-LRR genes played a critical role in resistance to FOL in tomato. Investigation of differentially regulated pathogen-induced NBS-LRR genes could lead to uncover the specific modulation patterns upon FOL infection in tomato.

To conclude, our abroad genome transcriptome RNA-seq data provided a comprehensive overview of the gene expression profiles between two different tomato cultivars Moneymaker and Motelle treated with FOL. Our results will facilitate further analysis of putative molecular mechanism of resistance in tomato upon to FOL, which eventually lead to improvement of Fusarium wilt disease resistance in tomato. It remains to be determined whether or how these candidate pathogen-related genes confirmed by qRT-PCR are overexpressed/knockouted in Moneymaker/Motelle plant to reveal the Fusarium wilt disease resistance. In this scenario, we would expect that overexpressing of these candidate pathogen-related genes will enhance resistance to *F. oxysporum* and would therefore develop a useful molecular tool to uncover functional roles for the increasing number of discovered genes in tomato.

## ACKNOWLEDGMENTS

We gratefully acknowledge support from JSSF: BK20161330, Jiangsu Province, China.

## LITERATURE CITED

Andolfo, G., F. Ferriello, L. Tardella, A. Ferrarini, L. Sigillo, et al., 2014 Tomato genome-wide transcriptional responses to Fusarium wilt and Tomato Mosaic Virus PLoS One 9: e94963.

Boller, T., and G. Felix, 2009 A renaissance of elicitors: perception of microbe-associated molecular patterns and danger signals by pattern-recognition receptors. Annu. Rev. Plant Biol. 60: 379–406.

Catanzariti, A.M., G.T. Lim, and D.A. Jones, 2015 The tomato I-3 gene: a novel gene for resistance to Fusarium wilt disease. New Phytol. 207:106–118.

Chen, Y.C., C.L. Wong, F. Muzzi, I. Vlaardingerbroek, B.N. Kidd, et al., 2014 Root defense analysis against *Fusarium oxysporum* reveals new regulators to confer resistance. Sci Rep. 4: 5584.

Cui, H., K. Tsuda, and J.E. Parker, 2014 Effector-Triggered Immunity: From Pathogen Perception to Robust Defense. Annu. Rev. Plant Biol. 66: 487–511.

Dangl, J.L., and J.D.G. Jones, 2001 Plant pathogens and integrated defence responses to infection. Nature 411: 826–833.

Dangl, J.L., and J.D.G. Jones, 2001 Plant pathogens and integrated defense responses to infection. Nature 411: 826–833.

De Ilarduya, O.M., A.E. Moore, and I. Kaloshian, 2001 The tomato Rme1 locus is required for Mi-1-mediated resistance to root-knot nematodes and the potato aphid. Plant J 27: 417–425.

Di Pietro, A., and M.I. Roncero, 1998 Cloning, expression, and role in pathogenicity of pg1 encoding the major extracellular endopolygalacturonase of the vascular wilt pathogen *Fusarium oxysporum*. Mol Plant Microbe Interact 11: 91–98.

Di Pietro, A., M.P. Madrid, Z. Caracuel, J. Delgado-Jarana, and M.I.G. Roncero, 2003 *Fusarium oxysporum:* exploring the molecular arsenal of a vascular wilt fungus. Molecular Plant Pathology. 4: 315–325.

Gonzalez-Cendales, Y., A.M. Catanzariti, B. Baker, D.J. Mcgrath, and D.A. Jones, 2016 Identification of I-7 expands the repertoire of genes for resistance to Fusarium wilt in tomato to three resistance gene classes. Mol Plant Pathol. 17: 448–463.

Guo, L., L. Han, L. Yang, H. Zeng, D. Fan, et al., 2014 Genome and transcriptome analysis of the fungal pathogen *Fusarium oxysporum* f. sp. *cubense* causing banana vascular wilt disease. PLoS One 9: e95543.

Gupta, K.J., L.A. Mur, and Y. Brotman, 2014 Trichoderma asperelloides suppresses nitric oxide generation elicited by *Fusarium oxysporum* in Arabidopsis roots. Mol Plant Microbe Interact 27: 307–314.

Houterman, P.M., L. Ma, G. van Ooijen, M.J. de Vroomen, B.J. Cornelissen, et al., 2009 The effector protein Avr2 of the xylem-colonizing fungus *Fusarium oxysporum* activates the tomato resistance protein *1–2* intracellularly. Plant J 58: 970–978.

Kawabe, M., Y. Kobayashi, G. Okada, I. Yamaguchi, T. Teraoka et al., 2005 Three evolutionary lineages of tomato wilt pathogen, *Fusarium oxysporum* f. sp. *lycopersici,* based on sequences of IGS, MAT1, and pg1, are each composed of isolates of a single mating type and a single or closely related vegetative compatibility group. J. Gen. Plant Pathol. 71: 263–272.

Langmead, B., C. Trapnell, M. Pop, and S.L. Salzberg, 2009 Ultrafast and memory-efficient alignment of short DNA sequences to the human genome. Genome biology 10: R25.

Lee, S.K., M.Y. Song, Y.S. Seo, H.K. Kim, S. Ko, et al., 2009 Rice Pi5-mediated resistance to *Magnaporthe oryzae* requires the presence of two coiled-coilnucleotide-binding-leucine-rich repeat genes. Genetics 181: 1627–1638.

Leslie, J.F. and B.A. Summerell, 2006 The Fusarium Laboratory Manual. Ames, IA: Blackwell Publishing. 388 pages p.

Liu, F., J.B. Wu, R.L. Zhan, and X.C. Ou, 2016 Transcription Profiling Analysis of *Mango-Fusarium mangiferae* Interaction. Front Microbiol. 7: 1443.

Liu, N., J. Yang, X. Fu, L. Zhang, K. Tang, et al., 2015 Genome-wide identification and comparative analysis of grafting-responsive mRNA in watermelon grafted onto bottle gourd and squash rootstocks by high-throughput sequencing. Mol Genet Genomics 291: 621–33.

Loutre, C., T. Wicker, S. Travella, P. Galli, S. Scofield, et al., 2009 Two different CC-NBS-LRR genes are required for Lr10-mediated leaf rust resistance in tetraploid and hexaploid wheat. Plant J 60: 1043–1054.

Marone, D., M.A. Russo, G. Laidò, A.M. De Leonardis, and A.M. Mastrangelo, 2013 Plant nucleotide binding site-leucine-rich repeat (NBS-LRR) genes: active guardians in host defense responses. Int J Mol Sci. 14: 7302–7326.

Merico, D., R, Isserlin, O. Stueker, A. Emili, and G.D. Bader, 2010 Enrichment Map: A Network-Based Method for Gene-Set Enrichment Visualization and Interpretation. PLoS ONE 5: e13984.

Mithöfer, A., and W. Boland, 2012 Plant defense against herbivores: chemical aspects. Annual Review of Plant Biology 63: 431–50.

Mukhtar, M.S., A.R. Carvunis, M. Dreze, P. Epple, J. Steinbrenner, et al., 2011 Independently evolved virulence effectors converge onto hubs in a plant immune system network. Science 333: 596–601.

Narusaka, M., K. Shirasu, Y. Noutoshi, Y. Kubo, T. Shiraishi, et al., 2009 RRS1 and RPS4 provide a dual Resistance-gene system against fungal and bacterial pathogens. Plant J 60: 218–226.

Okuyama, Y., H. Kanzaki, A. Abe, K. Yoshida, M. Tamiru, et al., 2011 A multifaceted genomics approach allows the isolation of the rice Pia-blast resistance gene consisting of two adjacent NBS-LRR protein genes. Plant J 66: 467–479.

Ori, N., Y. Eshed, I. Paran, G. Presting, D. Aviv, et al., 1997 The I2C family from the wilt disease resistance locus I2 belongs to the nucleotide binding, leucine-rich repeat superfamily of plant resistance genes. Plant Cell 9: 521–532.

Ouyang, S., G. Park, H. Atamian, C. Han, J. Stajich, et al., 2014 MicroRNAs suppress NB domain genes in tomato that confer resistance to *Fusarium oxysporum*. PLoS Pathogens. 10: e1004464.

Park, Y.S., H.J. Min, S.H. Ryang. K.J. Oh, J.S. Cha, et al., 2003 Characterization of salicylic acid-induced genes in Chinese cabbage. Plant Cell Reports 21: 1027–1034.

Paul, J.Y., D.K. Becker, M.B. Dickman, R.M. Harding, H.K. Khanna, et al., 2011 Apoptosis-related genes confer resistance to Fusarium wilt in transgenic ‘Lady Finger’ bananas. Journal of Plant Biotechnology 9: 1141–1148.

Peart, J.R., P. Mestre, R. Lu, I. Malcuit, and D.C. Baulcombe, 2005 NRG1, a CC-NBLRR protein, together with N, a TIR-NB-LRR protein, mediates resistance against tobacco mosaic virus. Curr Biol. 15: 968–973.

Ros B, Tummler F, Wenzel G (2004) Analysis of differentially expressed genes in a susceptible and moderately resistant potato cultivar upon *Phytophthora infestans* infection. Molecular Plant Pathology 5: 191–201.

Ruepp, A., A, Zollner, D, Maier, K, Albermann, J, Hani, et al., 2004 The FunCat, a functional annotation scheme for systematic classification of proteins from whole genomes. Nucleic acids research 32: 5539–5545.

Shah, J, 2003 The salicylic acid loop in plant defense. Current Opinion in Plant Biology 6: 365–371.

Simons, G., J. Groenendijk, J. Wijbrandi, M. Reijans, J. Groenen, et al., 1998 Dissection of the Fusarium *I2* gene cluster in tomato reveals six homologs and one active gene copy. Plant Cell 10: 1055–1068.

Sinapidou, E., K. Williams, L. Nott, S. Bahkt, M. Tor, et al., 2004 Two TIR:NB:LRR genes are required to specify resistance to *Peronospora parasitica* isolate Cala2 in Arabidopsis. Plant J 38: 898–909.

Smoot, M., K. Ono, J, Ruscheinski, P, Wang, and T. Ideker, 2011 Cytoscape 2.8: new features for data integration and network visualization. Bioinformatics 27: 431–432.

Smyth, G.K., 2005 Limma: linear models for microarray data. In: Gentleman R, Carey V, editors. Bioinformatics and Computational Biology Solutions Using R and Bioconductor. New York: Springer 397–420.

Storey, J.D., and R. Tibshirani, 2003 Statistical significance for genome wide studies. Proceedings of the National Academy of Sciences 100: 9440–9445.

Takken, F. and M. Rep, 2010 The arms race between tomato and *Fusarium oxysporum.* Mol Plant Pathol. 11: 309–314.

Trapnell, C., A. Roberts, L. Goff, G. Pertea, D. Kim, et al., 2012 Differential gene and transcript expression analysis of RNA-seq experiments with TopHat and Cufflinks. Nature protocols 7: 562–578

Trapnell, C., B.A. Williams, G. Pertea, A. Mortazavi, G. Kwan, et al., 2010 Transcript assembly and quantification by RNA-Seq reveals unannotated transcripts and isoform switching during cell differentiation. Nature biotechnology 28: 511–515.

Trapnell, C., L. Pachter, and S.L. Salzberg, 2009 TopHat: discovering splice unctions with RNA-Seq. Bioinformatics (Oxford, England) 25: 1105–1111.

Van Ooijen, G., H.A. van den Burg, B.J. Cornelissen, and F.L. Takken, 2007 Structure and function of resistance proteins in solanaceous plants. Annu. Rev. Phytopathol. 45: 43–72.

Xing, M., H. Lv, J. Ma, D. Xu, H. Li, et al., 2016 Transcriptome profiling of Resistance to *Fusarium oxysporum* f. sp. *conglutinans* in cabbage (Brassica oleracea) Roots. PLoS One 11: e0148048.

Yang, L., and H. Huang, 2014 Roles of small RNAs in plant disease resistance. J. Integr. Plant Biol. 56: 962–970.

Yu, S.C., and Y.M. Zou, 2008 A co-dominant molecular marker of Fusarium wilt resistance gene *I-2* derived from gene sequence in tomato. Yi Chuan 30: 926–932.

